# Efficacy and safety of decompressive craniectomy with non-suture duraplasty in patients with traumatic brain injury

**DOI:** 10.1101/2020.04.20.050799

**Authors:** Tae Seok Jeong, Gi Taek Yee, Tae Gyu Lim, Woo Kyung Kim, Chan Jong Yoo

**Affiliations:** Department of Neurosurgery, Gachon University Gil Medical Center, Incheon, Korea

## Abstract

**Background:** Decompressive craniectomy is an important surgical treatment for patients with severe traumatic brain injury (TBI). Several reports have been published on the efficacy of non-watertight sutures in duraplasty performed in decompressive craniectomy. This study aims to evaluate the effectiveness of dura closure without sutures (non-suture duraplasty) in decompressive craniectomy for TBI.

**Methods:** One hundred and six patients were enrolled at a single trauma center between January 2017 and December 2018. We retrospectively collected the data and classified the patients into non-suture and suture duraplasty craniectomy groups. We compared the characteristics of patients and their intra/post- operative findings such as operative time, blood loss, imaging findings, complications, and Glasgow outcome scale.

**Results:** There were 37 patients in the non-suture group and 69 in the suture craniectomy group. There were no significant differences between the two groups with regard to general characteristics. The operative time was 205 min for the suture duraplasty group and 150 min for the non-suture duraplasty group, and that for the non-suture duraplasty group was significantly lesser (p=0.002). Blood loss was significantly lesser in the non-suture duraplasty group (1000 mL) than in the suture duraplasty group (1500 mL, p=0.028). There were no other significant differences.

**Conclusion:** Non-suture duraplasty involved shorter operative time and lesser blood loss when compared to suture duraplasty. Other complications and prognosis were similar in both groups. Therefore, it can be concluded that decompressive craniectomy with non-suture duraplasty is a safe and useful surgical technique in patients with TBI.

## Introduction

Decompressive craniectomy is a neurosurgical procedure used for lowering the increased intracranial pressure in patients with severe traumatic brain injury (TBI). However, there is no clearly established standard guideline for the technique of decompressive craniectomy. Guresir et al. [1] reported that rapid closure decompressive craniectomy without duraplasty was a safe and feasible method for the management of malignant brain swelling. The technique of dura closure is mostly dependent on clinician experience. Dura suturing technique is traditionally known to require watertight closure to prevent complications such as cerebrospinal fluid (CSF) leak and infection. Several studies have reported that watertight duraplasty should not be used in decompressive craniectomy because non-watertight duraplasty can reduce the operative time while the probability of complications remains the same [1-3]. In cases of TBI, there are few reports on duraplasty performed without watertight suturing. Therefore, the purpose of this study was to determine if non-watertight dura reconstruction contributes significantly to the success of decompressive craniectomy in TBI.

## Materials and methods

### Patient population

This retrospective study was approved by our institutional review board (GCIRB2020-088). The requirement for obtaining informed consent from the patients was waived because the study was based on the information collected as part of routine clinical care and medical record-keeping. We reviewed the data of 151 TBI patients who underwent decompressive craniectomy and were hospitalized through a single trauma center between January 2017 and December 2018. We excluded patients who had previously undergone any surgical treatment for other brain lesions, patients younger than 18 years, and those over 70 years. We also excluded cases where the craniectomy was performed bilaterally or occipitally (posterior fossa), or if an emergency operation for another part of the body was simultaneously performed. As a result, 106 patients who underwent decompressive craniectomy due to TBI were enrolled in this study and divided into suture duraplasty and non-suture duraplasty groups.

### Data analysis

Data on general characteristics such as sex and age, and those pertaining to trauma such as Glasgow coma scale (GCS), anisocoria, and injury severity score (ISS) were reviewed retrospectively. We also reviewed intra/post-operative findings such as operative time, blood loss, and computed tomography (CT) scan, complications such as wound dehiscence and surgical site infection (SSI), and prognostic indicators such as the Glasgow outcome scale (GOS). The SSI was defined based on the US Centers for Disease Control and Prevention (CDC) criteria [4, 5].

### Surgical procedures

The standard decompressive craniectomy technique includes bone removal, dural incision, and hematoma removal. The dural incision was made in a star shape (Fig 1). Duraplasty was conducted in two manners: with and without suture. Suture duraplasty was done with watertight suturing of the dura with the artificial dura. Non-suture duraplasty was performed by overlaying an artificial dura over the dural opening area, without any suturing (Fig 2). The choice of surgical technique (suture or non-suture duraplasty) depended on the surgeon’s preference. After duraplasty, the subcutaneous tissue and skin were closed in a serial fashion.

**Fig 1.**
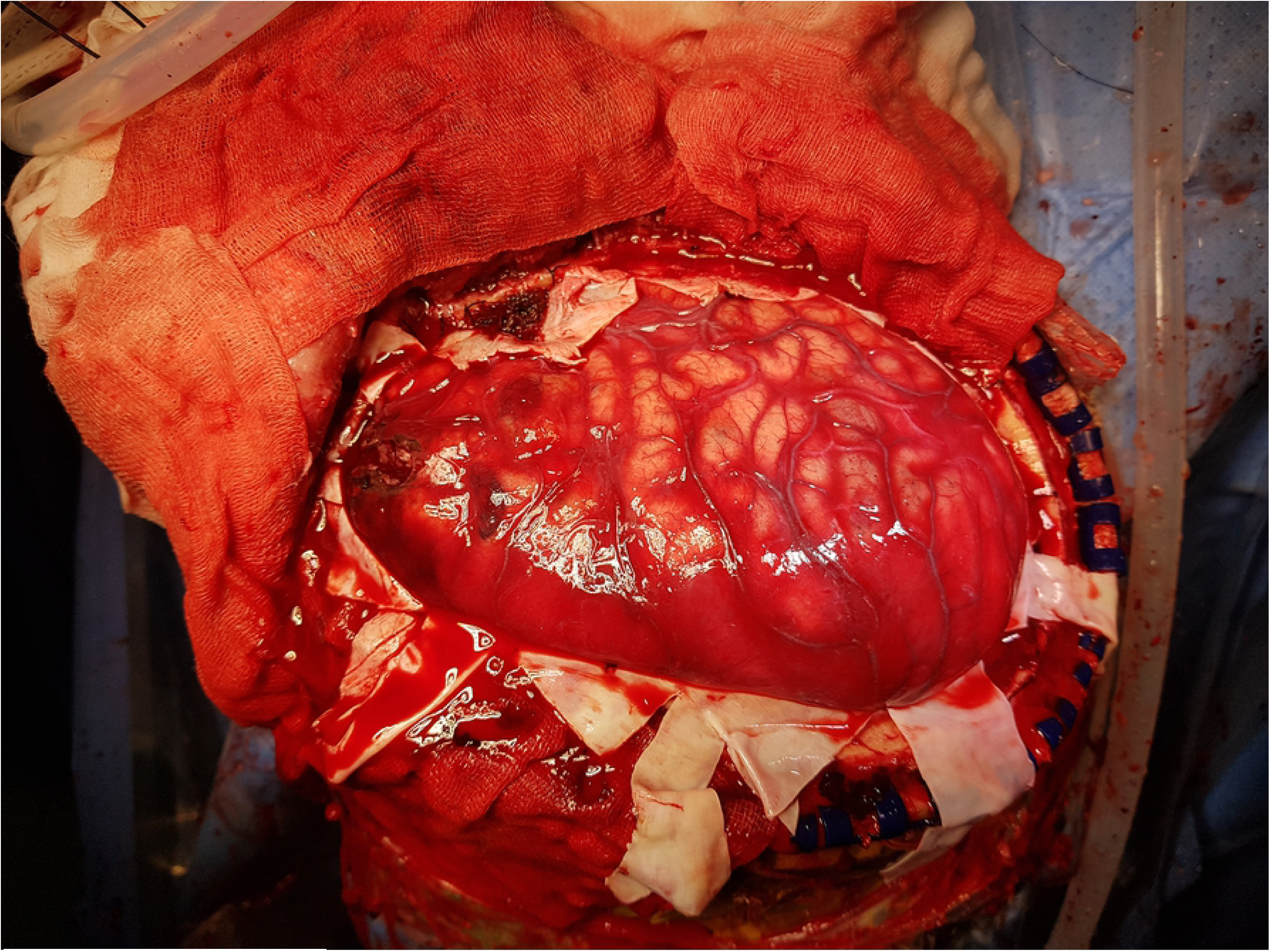
Intraoperative photograph showing wide craniotomy and dura opening.

**Fig 2.**
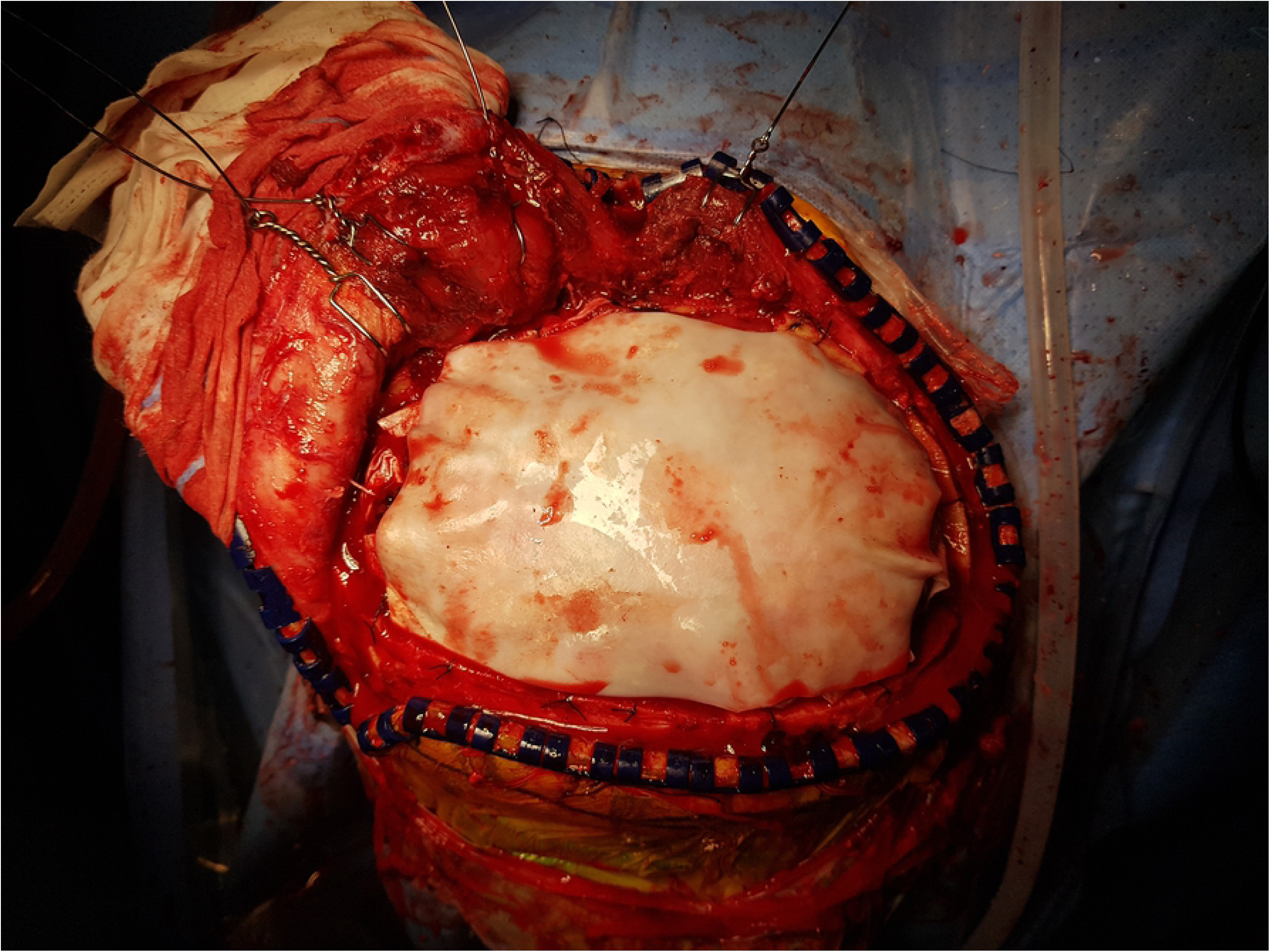
Intraoperative photograph showing non-suture duraplasty with an artificial dura material overlaid on the brain cortex.

### Statistical analysis

Continuous data (e.g., age) were presented as medians and interquartile ranges. Categorical data (e.g., sex) were presented using frequencies and percentages. Continuous variables were compared using the independent t-test and categorical variables were compared using the Fisher exact or Pearson chi-square test. All tests were performed using a statistical significance criterion of α=0.05, and analyses were performed in SPSS for Windows version 23.0 (IBM Corp., Armonk, NY, USA).

## Results

### Patient characteristics

Of the 106 patients (female: 31, male: 75), 69 belonged to the suture duraplasty group and 37 to the non-suture duraplasty group. The median age of the subjects was 53 years, and the most common diagnosis given was subdural hematoma (76.4%). The GCS score of the subjects at admission was classified into three categories according to TBI severity: mild (GCS: 13∼15), moderate (9∼12), and severe (3∼8). Severe TBI (67.9%) was the most common. Median ISS was 25. There were no significant differences between the two groups with respect to patient characteristics (Table 1).

**Table 1.** Patient characteristics. GCS: Glasgow Coma Scale; ISS: Injury Severity Score. Values are presented as median value

|  | **Suture**<br>**duraplasty**<br>**(n=69)** | **Non-suture**<br>**duraplasty**<br>**(n=37)** | **Total**<br>**(n=106)** | ***p*-value** |
| --- | --- | --- | --- | --- |
| Sex |  |  |  | 0.597 |
| Female | 19 (27.5) | 12 (32.4) | 31 (29.2) |  |
| Male | 50 (72.5) | 25 (67.6) | 75 (70.8) |  |
| Age (years) | 52 (42-60) | 56 (43-59) | 53 (42-60) | 0.942 |
| Diagnosis |  |  |  | 0.269 |
| Subdural hematoma | 55 (79.7) | 26 (70.3) | 81 (76.4) |  |
| Epidural hematoma | 7 (10.1) | 8 (21.6) | 15 (14.2) |  |
| Intracerebral hematoma | 7 (10.1) | 3 (8.1) | 10 (9.4) |  |
| GCS score |  |  |  | 0.723 |
| 3-8 | 48 (69.6) | 24 (64.9) | 72 (67.9) |  |
| 9-12 | 9 (13.0) | 7 (18.9) | 16 (15.1) |  |
| 13-15 | 12 (17.4) | 6 (16.2) | 18 (17.0) |  |
| Anisocoria |  |  |  | 0.853 |
| Yes | 33 (47.8) | 17 (45.9) | 50 (47.2) |  |
| No | 36 (52.2) | 20 (54.1) | 56 (52.8) |  |
| ISS | 25 (17-30) | 25 (24-33) | 25 (21-33) | 0.152 |

(interquartile ranges) or number (%).

### Intra/post-operative findings

As intraoperative findings, operative time and blood loss were analyzed. The operative time was 205 min for the suture duraplasty group and 150 min for the non-suture duraplasty group; the non-suture duraplasty group had significantly shorter operative time (p=0.002). In addition, blood loss was significantly lesser in the non-suture duraplasty group (1000 mL) than in the suture duraplasty group (1500 mL, p=0.028).

A postoperative brain CT scan was done one month after surgery. The amount of subgaleal fluid collection did not differ between the two groups, and ipsilateral/contralateral subdural hygroma tended to be more in the suture duraplasty group; however, there was no statistically significant difference. Wound dehiscence and SSI were evaluated for the complication, and there were no significant differences between the two groups. GOS score at discharge also did not differ between the two groups (Table 2).

**Table 2.** Intra/post-operative findings and results.

|  | **Suture duraplasty**<br>**(n=69)** | **Non-suture duraplasty**<br>**(n=37)** | ***p*-value** |
| --- | --- | --- | --- |
| Intraoperative findings |  |  |  |
| Operative time (min) | 205 (160-265) | 150 (105-225) | 0.002* |
| Blood loss (mL) | 1500 (1000-2500) | 1000 (700-1800) | 0.028* |
| Postoperative CT findings |  |  |  |
| Subgaleal fluid collection | 12 (17.4) | 9 (24.3) | 0.648 |
| Ipsilateral subdural hygroma | 16 (23.2) | 4 (10.8) | 0.121 |
| Contralateral subdural hygroma | 12 (17.4) | 2 (5.4) | 0.131 |
| Complications |  |  |  |
| Wound dehiscence | 10 (14.5) | 4 (10.8) | 0.766 |
| Surgical site infection | 6 (8.7) | 2 (5.4) | 0.710 |
| GOS score |  |  | 0.834 |
| 1 | 9 (13.0) | 6 (16.2) |  |
| 2 | 12 (17.4) | 9 (24.3) |  |
| 3 | 25 (36.2) | 13 (35.1) |  |
| 4 | 17 (24.6) | 7 (18.9) |  |
| 5 | 6 (8.7) | 2 (5.4) |  |

CT: Computed Tomography; GOS: Glasgow Outcome Scale. Values are presented as median value (interquartile ranges) or number (%). \*Statistically significant differences (*p*-value <0.05).

## Discussion

It is normal for individual differences to exist in the method of dura reconstruction between surgeons, and they vary based on the training received. Traditionally, many surgeons recommend tightly closing the dura mater (i.e., ‘watertight suture’) because this has been thought to prevent various complications such as infection, CSF leakage, and subgaleal fluid collection. However, the efficacy of watertight suturing is not entirely known, especially in TBI cases. Recently, some authors have argued that watertight suturing increases operative time and hospital costs, and the occurrence of complications is similar to that in cases without watertight sutures [1, 2, 6]. Guresir et al.[1] introduced rapid closure decompressive craniectomy, which is not dura reconstruction, but filling of the dural defect with a hemostat material. Non-suture duraplasty craniectomy is similar to rapid closure craniectomy, but uses synthetic dura instead of the hemostat material to avoid adhesion in cranioplasty. This difference may create higher costs for non-suture duraplasty craniectomy than rapid closure craniectomy, although there may be surgical benefits such as shortened operative time due to inhibited adhesion at cranioplasty subsequent to non-suture duraplasty craniectomy.

The present study evaluated the operative time and blood loss as the comparable intra-operative parameters. The non-suture duraplasty craniectomy group revealed shorter operative time and lesser blood loss than the suture group. It is known that shorter operative time and lower bleeding levels aid in decreasing the possibility of infection and hospital costs [7, 8]. It also may help treat patients requiring critical care, such as multiple trauma [9, 10].

The incidence of postoperative wound infection documented in the literature ranges from as low as 1.25% to as high as 17% without, and 0.3% to 3.0% with prophylactic antibiotics [9-13]. The SSI incidence in the present study was approximately 7.5%, which is slightly higher than that in previous studies. Korinek et al. [9] reported that independent predictive risk factors of SSI after craniotomy included recent neurosurgery, contaminated wounds, prolonged operative time, and emergency operation. A slightly higher infection rate in the present study was attributed to the fact that all patients included in the study underwent emergency surgery. In the present study, the SSI incidence of the non-suture duraplasty craniectomy group was lower than that of the suture group, although there was no statistical significance. This may be because the operative time of the non-suture duraplasty craniectomy group was shorter than that of the suture group.

The incidence of subdural hygroma after decompressive craniectomy in TBI patients was reported to be 21-50% [6, 14-17]. Subdural hygroma frequently occurs on the ipsilateral side of the craniectomy; however, it can occur on the contralateral side or bilaterally. Surgical treatment is not always required, but may be necessary if it converts to a chronic subdural hematoma or is caused by a mass effect [15, 16]. In the present study, the incidence of subdural hygroma (both ipsilateral and contralateral) was about 32.1% (similar to previous studies), and was lesser in the non-suture duraplasty craniectomy group than the suture group. If watertight duraplasty is performed, even in conjunction with decompressive duraplasty, the brain may still be compressed by the dura or subdural fluid because the progress of brain edema cannot be predicted. In cases of insufficient decompressive duraplasty, subdural hygroma is more likely to occur due to a decrease in the infusion of CSF circulation and brain shift [6]. Therefore, non-suture duraplasty craniectomy may help to reduce the incidence of subdural hygroma, even in cases of severe brain edema, because there is no compression from the dura. However, the above results are from preliminary reports with insufficient statistical evidence.

Subgaleal fluid collection was somewhat higher in the non-suture duraplasty craniectomy group. Vieira et al. [3] reported that once the arachnoid is intact, there is no increased risk of CSF leaks, and watertight closure may lead to small defects on suture lines, causing a “one-way valve” effect that could potentially facilitate the development of CSF leakage. In this study, there was no analysis of arachnoid injury, so this could not be accurately evaluated. However, we think that the presence of subgaleal fluid collection is not meaningful because there were no differences in postoperative complications and GOS.

We observed 37 patients who received cranioplasty following decompressive craniectomy. GOS scores did not differ between the two groups (suture vs. non-suture), and patients with severe head injuries had poorer prognosis. Therefore, we do not believe that watertight suturing necessarily leads to good prognosis. In the non-suture duraplasty craniectomy group, the brain cortex was not exposed because of the formation of fibrotic tissue between the scalp flap and dura material (Fig 3). The incidence of complications, including infection after cranioplasty, were similar in the suture duraplasty and non-suture duraplasty groups (3% and 4%, respectively). Therefore, we believe that non-suture duraplasty may not cause any problems in cranioplasty, although further evaluation is needed.

**Fig 3.**
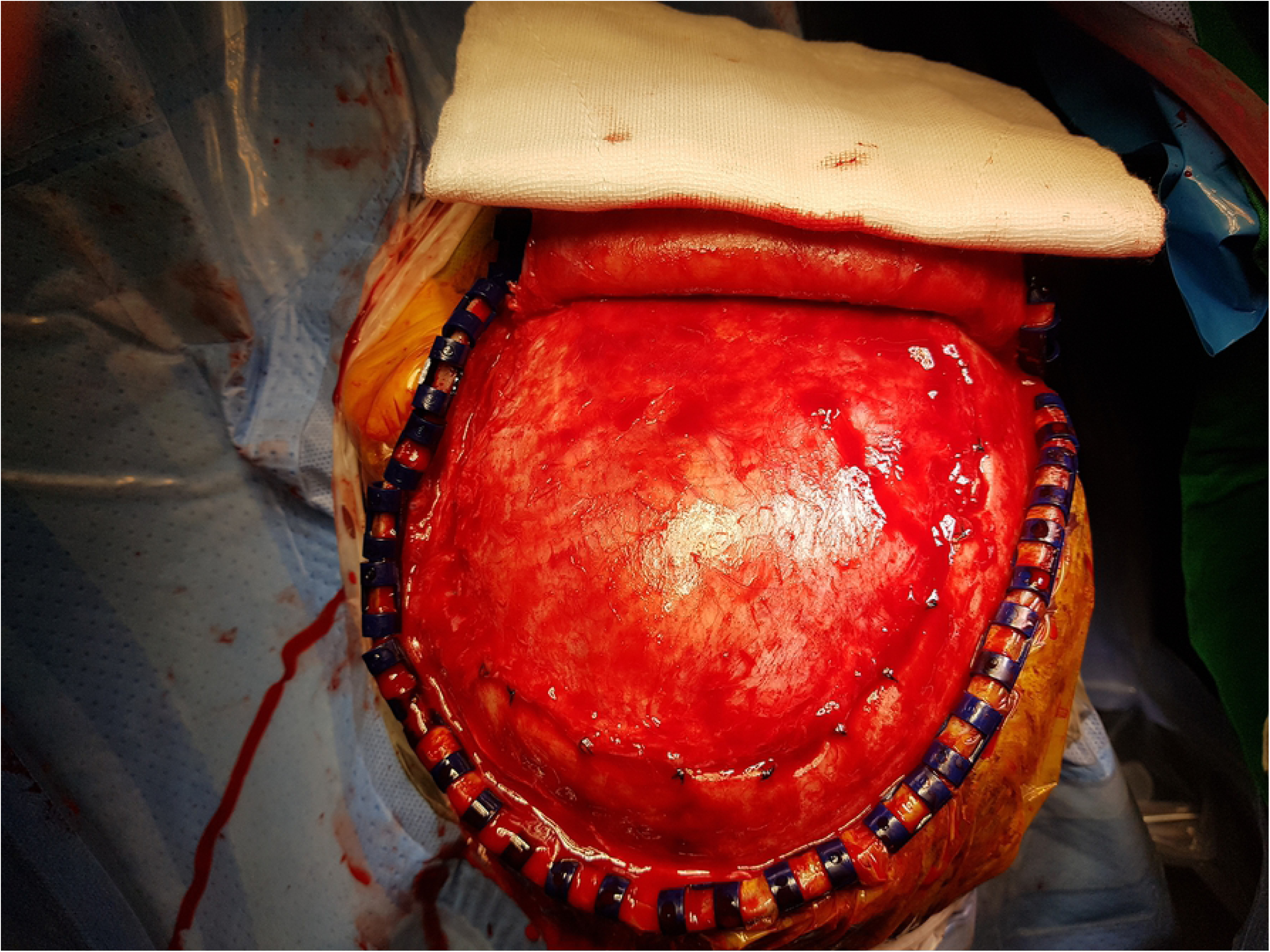
Intraoperative photograph showing craniectomy site covered by fibrotic tissue in cranioplasty following decompressive craniectomy with non-suture duraplasty.

This study has some limitations. The data were analyzed retrospectively and the sample size was relatively small. Also, a decompressive craniectomy was performed by multiple surgeons. It may influence intra/post-operative results such as operative time, blood loss, and prognosis. Despite these limitations, the results of this study may provide useful information regarding the benefits of non-suture duraplasty, such as a shorter operative time and lesser blood loss during decompressive craniectomy.

## Conclusions

This study is a report on non-suture duroplasty performed as a part of decompressive craniectomy in TBI patients. Non-suture duraplasty revealed a shorter operative time and lesser blood loss when compared to suture duraplasty. Other complications and prognosis were similar in both groups. Therefore, it can be assumed that decompressive craniectomy with non-suture duraplasty is safe and effective in patients with TBI.

## Acknowledgments

We thank Dae Han Choi for the technical work on graphics.

